# Genome-scale CRISPR‒Cas9 screen identifies novel host factors as potential therapeutic targets for SARS-CoV-2 infection

**DOI:** 10.1101/2023.03.06.531431

**Authors:** Madoka Sakai, Yoshie Masuda, Yusuke Tarumoto, Naoyuki Aihara, Yugo Tsunoda, Michiko Iwata, Yumiko Kamiya, Ryo Komorizono, Takeshi Noda, Kosuke Yusa, Keizo Tomonaga, Akiko Makino

**Author notes:** Corresponding author: Akiko Makino.

## Abstract

Although many host factors important for severe acute respiratory syndrome coronavirus 2 (SARS-CoV-2) infection have been reported, the mechanisms by which the virus interacts with host cells remain elusive. Here, we identified tripartite motif containing (TRIM) 28, TRIM33, euchromatic histone lysine methyltransferase (EHMT) 1, and EHMT2 as novel proviral factors involved in SARS-CoV-2 infection by CRISPR‒Cas9 screening. We demonstrated that TRIM28 plays a role(s) in viral particle formation and that TRIM33, EHMT1, and EHMT2 are involved in viral transcription and replication using cells with suppressed gene expression. UNC0642, a compound that specifically inhibits the methyltransferase activity of EHMT1/2, strikingly suppressed SARS-CoV-2 growth in cultured cells and reduced disease severity in a hamster infection model. This study suggests that EHMT1/2 may be a novel therapeutic target for SARS-CoV-2 infection.

## Introduction

Therapeutics that target proviral factors have the potential to be effective against a broad spectrum of viruses. Genome-wide screens using CRISPR‒Cas9 technology to identify host factors involved in SARS-CoV-2 replication have been performed in several studies (1–12). The only host factor identified in every screen is ACE2, and different genes are identified depending on conditions such as the cells used, the multiplicity of infection (MOI) and the time to sample collection (4, 13).

A549 cells, which are human lung epithelial type II cells, typically do not show clear cytopathic effects from SARS-CoV-2 infection. In previous studies, proviral factors were found by prolonging the sampling time (3) or by using a mutant virus lacking the S1/S2 sites (1). We discovered that A549-hACE2 cells infected with purified SARS-CoV-2 caused clear cell death and applied this to our CRISPR‒Cas9 screen.

In this study, we successfully identified four new genes from the same family or complex as proviral factors of SARS-CoV-2. TRIM28 and TRIM33 are members of the same gene family (14) but have different effects on in the virus life cycle. EHMT1/2 form heterodimers (15, 16), and depletion of their expression suppresses viral proliferation. A specific inhibitor of EHMT1/2 suppressed viral growth *in vitro* and *in vivo*. This study contributes to a new understanding of how SARS-CoV-2 interacts with host cells and to the development of novel methods to control the virus by targeting proviral factors.

## Results

### Identification of novel proviral factors for SARS-CoV-2 infection

We transduced a single guide RNA (sgRNA) library targeting 18,365 human genes with 113,526 sgRNAs, including 1,004 nontargeting control gRNAs (17, 18), into A549-hACE2-Cas9 cells and then infected them with purified SARS-CoV-2 UT-NCGM02 (19) at an MOI of 0.3 on day 10 posttransduction. DNA was harvested from sgRNA-transduced cells at 10 days posttransduction and from the mock or infected cells at 3 days postinfection for NGS analysis (Fig. 1a). By comparing the number of reads of sgRNA in the day 10 sample with mock or infected cells, we found that the control gRNA (gCTRL) was linearly aligned, indicating that there was no bias due to the experimental technique (Supplementary Data 1a). In addition to *ACE2*, *VPS35*, and *VPS29,* which have been previously reported (1, 3, 20), we detected *MED12, MED23, EHMT1, EHMT2, RPL34,* and *TRIM28* as target genes enriched in infected cells that showed resistance to infection-induced cell death by plotting the comparison of sgRNA read counts between the two groups based on log-fold change or statistic score. Conversely, *DAXX, NF2, DAZAP2*, and *TRIM33* were enriched in the mock cells (Fig. 1b). In the comparison of mock and infected cells, *MED12* and *TRIM28* were positively selected in addition to *ACE2*, whereas *NF2* was under strong negative selection (Supplementary Data. 1b).

**Fig. 1.**
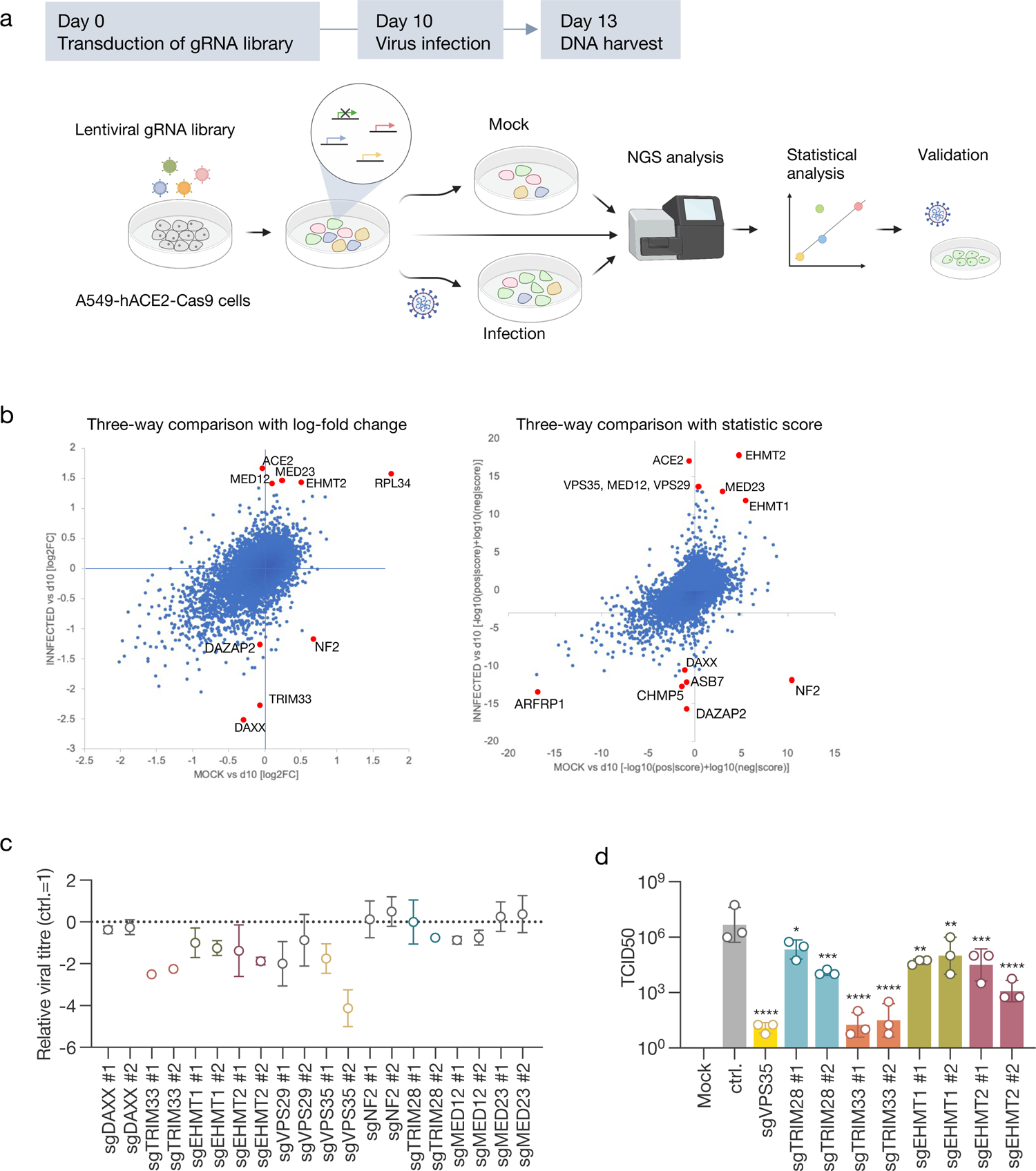
Genome-wide CRISPR‒Cas9 screening to identify host factors for SARS-CoV-2 infection. **(a)** Schematic diagram of the screening. ACE2- and Cas9-expressing A549 cells transduced with a lentiviral-packaged whole-genome sgRNA library were infected with purified SARS-CoV-2. DNA was extracted from surviving cells 3 days postinfection and analysed for gRNA. **(b)** Scatterplot of the results from screening. Comparison of gRNA changes between preinfection (d10) versus mock and preinfection (d10) versus infected. **(c)** Comparison of viral titres in target gene knockout cells. The knockout cells were infected with SARS-CoV-2. After 3 days, the supernatant was harvested and the viral titre was evaluated by TCID50. The titre collected from control cells was used as a standard. **(d)** Titration of viral production in the selected gene knockout cells for validation. The titre was determined by TCID50 as in (c). Data are presented as the mean ± the standard deviation (SD) of three independent experiments. Statistical analysis was performed using Fisher’s least significant difference (LSD) test. *, *p* < 0.05; **, *p* < 0.01; ***, *p* < 0.001; ****, *p* < 0.0001.

Interestingly, genes that form families or complexes were identified as host factors associated with SARS-CoV-2 infection. As a validation experiment, we generated individual knockout cells by transducing A549-hACE2-Cas9 cells with sgRNA. The knockout cells were infected with UT-NCGM02 at an MOI of 0.001 and then the virus titre in the supernatant was measured 3 days later. The viral titre was reduced in cells transduced with sgRNA targeting *TRIM28*, *TRIM33*, *EHMT1*, and *EHMT2* (Fig 1c). In cells with low knockout efficiency, the inhibitory effect on virus replication was also low (Fig. 1d, Supplementary Data 2a, b). The viral titre was not reduced in the cells transduced with sgRNA targeting *MED12*, *MED23*, *NF2*, and *DAXX* (Fig. 1c).

### TRIM28 and TRIM33 are involved in the replication of SARS-CoV-2 through different mechanisms

Since TRIM28 and TRIM33 have been reported to negatively regulate the production of interferon β (IFNβ) (21–23), we quantified the amount of *IFNB1* mRNA in A549-hACE2-Cas9 cells transduced with sgRNA targeting *TRIM28* (sgTRIM33 cells) or *TRIM33* (sgTRIM28 cells). The results did not show upregulation of *IFNB1* mRNA in either sgTRIM28 or sgTRIM33 cells (Fig. 2a), suggesting that SARS-CoV-2 replication is not suppressed by promoting IFNβ production in the cells. To evaluate how the identified proviral factors are involved in the replication of SARS-CoV-2, we then infected sgTRIM28 and sgTRIM33 cells with UT-NGGM02 and quantified the expression of viral RNA and protein in the cells and supernatants. Compared to sgCTRL cells, sgVPS35 and sgTRIM33 cells exhibited downregulated expression of viral RNA and protein in both cell lysates and supernatants, while it was upregulated in sgTRIM28-induced cells (Fig. 2b, c). Furthermore, SARS-CoV-2 spike-mediated infection efficiency was not reduced in sgTRIM28- and sgTRIM33-induced cells in a pseudotyping assay using vesicular stomatitis virus (VSV)ΔG (Fig. 2d). In sgCTRL cells, the SARS-CoV-2 spike protein was localized to the ER-Golgi intermediate compartment, where viral assembly occurs (24–27). However, in sgTRIM28 cells, it was widely distributed (Fig. 2e). Electron microscopy analysis also showed that no viral particles were produced in the sgTRIM28 cells (Fig. 2f). These results suggest that TRIM28 and TRIM33 are not involved in the entry step of SARS-CoV-2 and that TRIM28 may play a role in the formation of virus particles, while TRIM33 may be involved in the transcription and/or replication process in the cell.

**Fig. 2.**
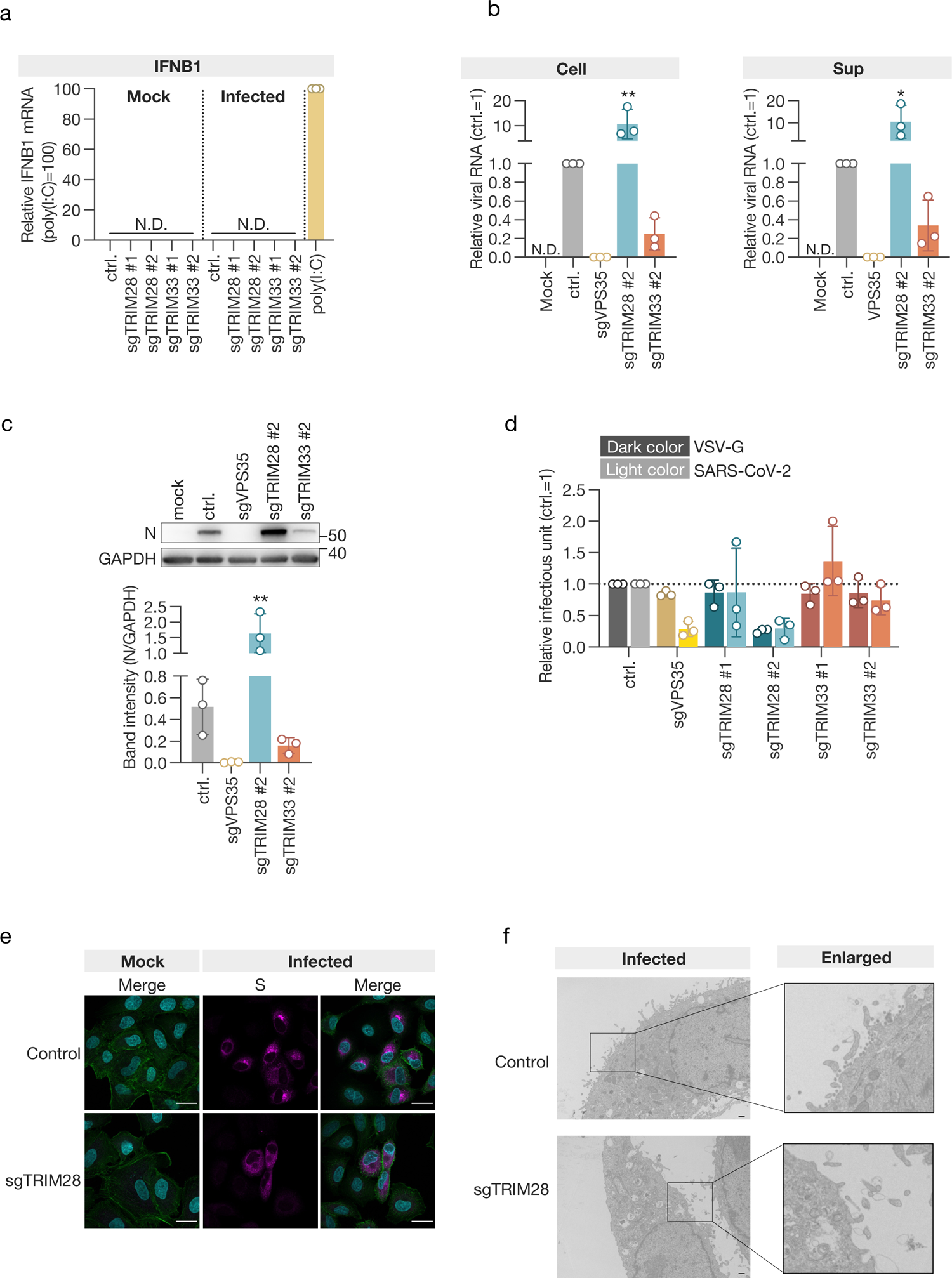
TRIM28 and TRIM33 are proviral factors for SARS-CoV-2 infection. **(a)** Quantification of *IFNβ* mRNA at 24 hours post-infection. Poly:IC (HMW) was transfected as a positive control. **(b)** qPCR analysis of viral RNA. **(c)** Western blotting analysis of the viral nucleoprotein (N). The expression of viral protein was evaluated by the band intensity of N, which was normalized to that of GAPDH. Cell lysates (b, c) and culture supernatants (b) were harvested 3 days after infection. **(d)** VSV-based pseudotype assay. VSVΔG-GFP enveloped with VSV-G or SARS-CoV-2-S was used to infect knockout cells, and the number of GFP-positive cells was counted. **(e)** The subcellular localization of the spike protein in the infected cells. Cells were stained with anti-S antibody and phalloidin for visualization of cellular actin. Bar = 25 μm. **(f)** Electron microscopic analysis of virus particle formation in infected cells. Bar=5.0 μm. Data are presented as the mean ± the SD of three independent experiments. Statistical analysis was performed using Fisher’s LSD test. *, *p* < 0.05; **, *p* < 0.01. N.D., not detected.

### EHMT1/2 inhibitor suppresses the proliferation of SARS-CoV-2 *in vitro*

EHMT1 and EHMT2 (also referred to as GLP and G9a) form heterodimers in cells and methylate histone H3 as well as nonhistone proteins (15, 16, 28–30). EHMT1/2 is important for inhibiting IFNγ-induced chemokine transcription (31, 32), and in bovine cells, inhibition of EHMT2 increased IFNβ transcription (33). Therefore, we transduced A549-hACE2-Cas9 cells with sgRNAs targeting *EHMT1* and *EHMT2* (sgEHMT1 cells and sgEHMT2 cells, respectively) and quantified the mRNA levels of *IFNB1* and *CXCL10*. As shown in Fig. 3a, the expression inhibition of EHMT1 and EHMT2 in the cells we used did not cause an increase in *IFNB1* and *CXCL10* mRNA levels, suggesting that EHMT1 and EHMT2 do not suppress the proliferation of SARS-CoV-2 by inhibiting the production of IFNβ and cytokines. We then quantified viral RNA and protein in the cell lysate and supernatant of SARS-CoV-2-infected cells and found that both viral RNA and protein expression were downregulated in sgEHMT2 cells, while no differences were observed in sgEHMT1 cells compared to sgCTRL cells (Fig. 3b, c). Similarly, using a VSV-based pseudotype assay, the spike entry efficiency of SARS-CoV-2 was not reduced in sgEHMT1 and sgEHMT2 cells (Fig. 3d).

**Fig. 3.**
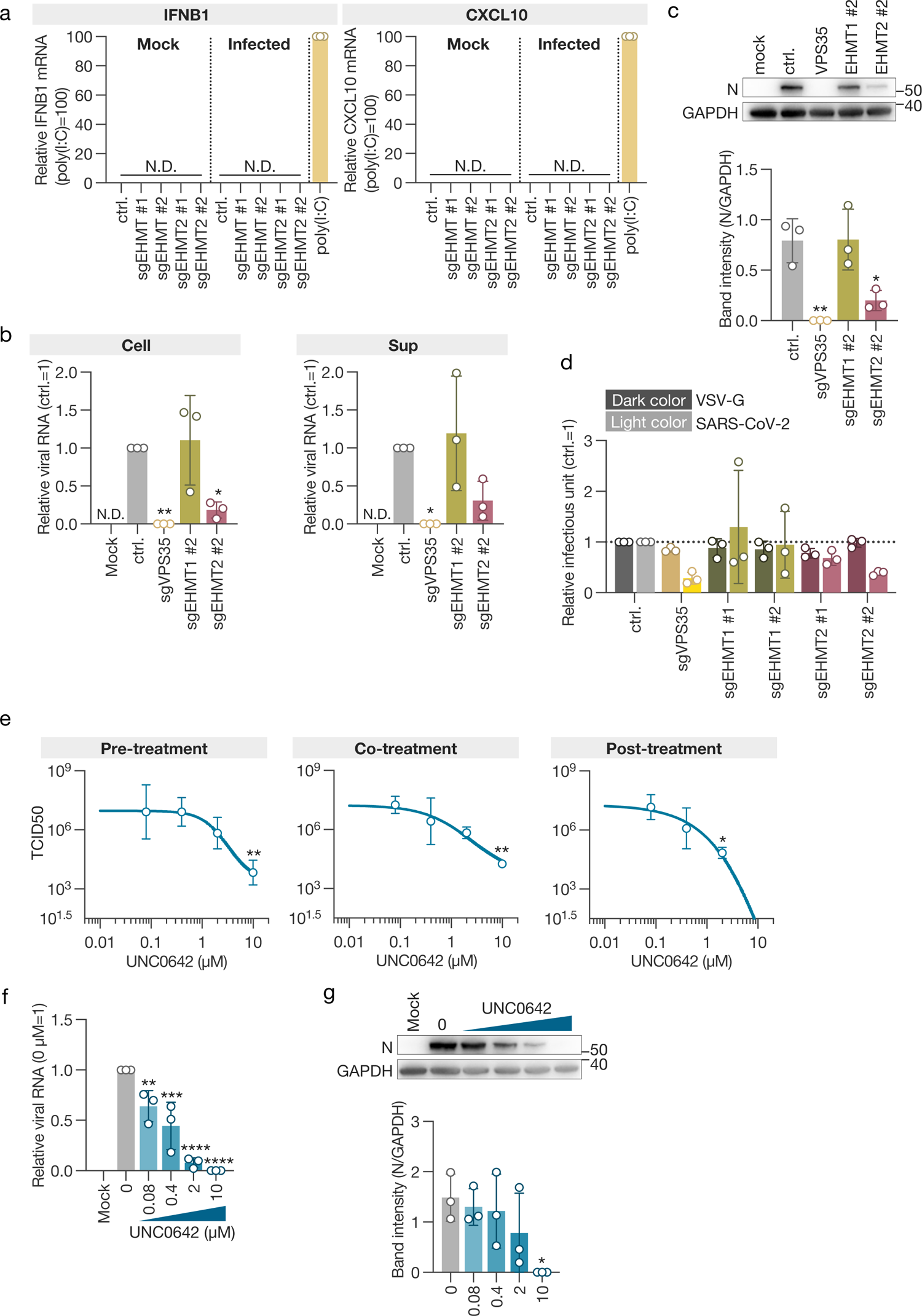
EHMT1 and EHMT2 are proviral factors for SARS-CoV-2 infection. **(a)** Quantification of mRNA for *IFNβ* and *CXCL10*. Poly:IC (HMW) was transfected as a positive control. **(b)** qPCR analysis of viral RNA. **(c)** Western blotting analysis of viral N protein. **(d)** VSV-based pseudotype assay. **(e)** Inhibitory effect of the EHMT1/2-specific inhibitor UNC0642 upon SARS-CoV-2 infection. **(f)** qPCR analysis of viral RNA in UNC0642-treated cells. **(g)** Western blotting analysis of viral N protein in UNC0642-treated cells. The expression of viral protein was evaluated by the band intensity of N, which was normalized to that of GAPDH. Data are presented as the mean ± the SD of three independent experiments. Statistical analysis was performed using Fisher’s LSD test (b, c, f, g) and Dunnett’s multiple-comparison test (e). *, *p* < 0.05; **, *p* < 0.01; ***, *p* < 0.001; ****, *p* < 0.0001.

UNC0642 is a methyltransferase inhibitor with high selectivity for EHMT1/2 and low cytotoxicity (Supplementary Data 3a) (34, 35). To evaluate whether the enzymatic activity of EHMT1/2 is involved in the proliferation of SARS-CoV-2, we treated the cells with the inhibitor before, during and after virus inoculation and evaluated the virus titre in the supernatant. The results showed that remdesivir, which inhibits viral RNA-dependent RNA polymerase (36, 37), suppressed virus proliferation only after inoculation (Supplementary Data. 3b), whereas UNC0642 consistently and dose-dependently suppressed SARS-CoV-2 proliferation under all conditions (Fig. 3e). Specifically, when we treated with 10 μM UNC0642 after virus infection, virus proliferation was suppressed by approximately 10^6^ (Fig. 3e). When treated with UNC0642 after virus infection, the expression of virus RNA and protein in the cells was also suppressed in a dose-dependent manner (Fig. 3f, g). These results demonstrate that EHMT1/2, through its enzymatic activity, is involved in the transcription and/or replication of SARS-CoV-2 in the cell.

### EHMT1/2 inhibitor reduces SARS-CoV-2 growth and disease severity in infected animals

To assess whether UNC0642 also suppresses the growth of SARS-CoV-2 *in vivo*, we inoculated 4-week-old male Syrian hamsters with the virus and administered the inhibitor at a dose of 5 mg/kg once a day intraperitoneally (Fig. 4a). On day 4 postinoculation, the lungs of the animals were subjected to virus titration and pathological analysis. The UNC0642-treated group lost weight due to viral infection, as did the DMSO-treated group, but recovery after 4 days of infection was higher than that in the DMSO-treated group (Fig. 4b). Viral growth in the lungs was suppressed in the UNC0642-treated group compared to the DMSO-treated group (Fig. 4c). Using a semiquantitative pathological scoring system (38, 39) to evaluate the infected animals, the cumulative score was significantly lower in the UNC0642-treated group than in the DMSO-treated group (Fig. 4d, e). In particular, the scores for vessels and regeneration were lower in the UNC0642-treated group (Supplementary Data 4). These results demonstrate that the selective inhibitor of EHMT1/2, UNC0642, also suppresses the growth of SARS-CoV-2 and reduces the severity of infection *in vivo*.

**Fig. 4.**
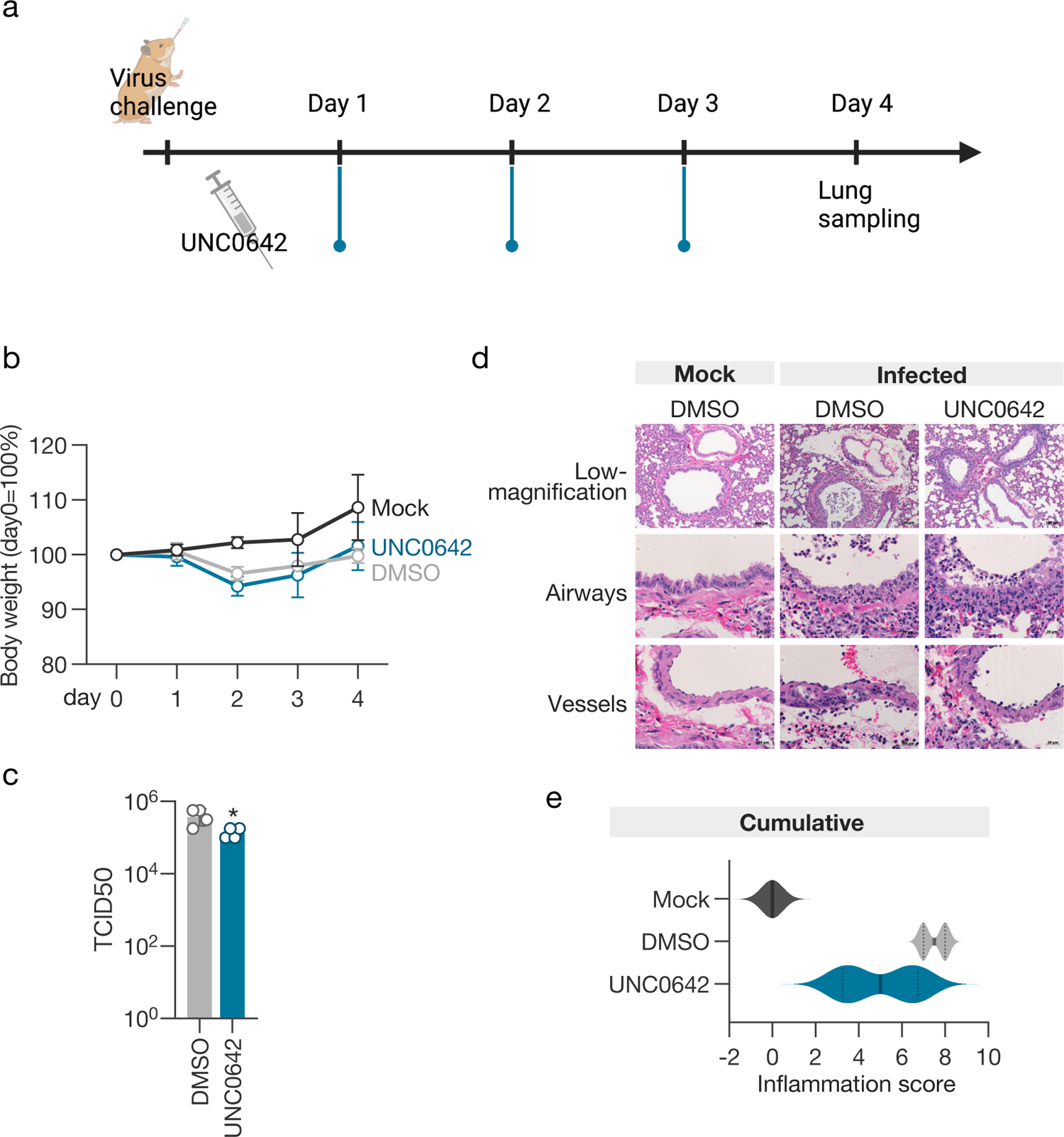
EHMT1/2 inhibitor reduces SARS-CoV-2 growth and disease severity in infected animals. **(a)** Timeline of virus challenge and inhibitor administration. **(b)** Viral load in the lungs of hamsters 4 days post-infection. **(c)** Body weight change of hamsters. **(d)** Representative histopathological images of the lungs of hamsters. **(e)** Violin plot of the cumulative inflammation score. The data are presented as the mean ± the SD of three independent experiments. Statistical analysis was performed using an unpaired t test. *, *p* < 0.05.

## Discussion

We used CRISPR‒Cas9-mediated genome-wide screening to identify TRIM23, TRIM33, and EHMT1/2 as proviral factors of SARS-CoV-2. Using purified virus, we were able to perform a genetic screen using cell death as an indicator, which allowed us to identify previously unreported proviral factors. This screening also extracted DAXX, which has been reported as a restriction factor (12). However, when we transduced sgRNA into A549-hACE2-Cas9 cells and assessed virus replication, we did not observe any promotion of virus replication (Fig. 1C). This may be due to differences in knockout efficiency or cell types used (Supplementary Data 2).

TRIM28 and TRIM33 are E3 ubiquitin ligases involved in the regulation of innate immunity (21–23, 40). In A549-hACE2-Cas9 cells in which TRIM28 or TRIM33 was knocked out, we did not observe any increase in *IFNB1* mRNA expression levels (Fig. 2a). In addition, it has been reported that suppression of EHMT2 expression in bovine cells leads to an increase in IFNβ production (33), but in sgEHMT1 or EHMT2 cells, there was no increase in *IFNB1* mRNA levels (Fig. 3a). These results suggest that TRIM28, TRIM33, and EHMT1/2 do not promote SARS-CoV-2 proliferation by suppressing the antiviral response but are involved in the viral growth cycle. TRIM33 and TRIM28 are members of the same TRIM family but are involved in SARS-CoV-2 replication through different mechanisms (Fig. 2). Several SARS-CoV-2 proteins that are ubiquitinated have been reported (41–45), but further analysis is needed to determine whether TRIM28 and TRIM33 are involved in the ubiquitination of viral proteins.

EHMT1/2 catalyses the methylation of histones and nonhistone proteins, and the substrates of the enzymes that contribute to SARS-CoV-2 replication may be either host or viral proteins or both. There are reports that EHMT2 is involved in numerous gene pathways that promote translation, many of which are also involved in SARS-CoV-2 replication (46, 47). Moreover, ORF8 of SARS-CoV-2 mimics histones through the ARKS motif and disrupts epigenetic control in infected cells (48). Furthermore, inhibitors that block viral replication, whether applied before or after infection (Fig. 3e), suggest that EHMT1/2 is involved in SARS-CoV-2 replication in a complex manner.

The development of host-targeted antiviral drugs has not been active due to potential side effects; however, there is a growing need for a variety of strategies to control viral infections. In this study, UNC0642, which selectively inhibits the enzymatic activity of EHMT1/2, significantly suppressed SARS-CoV-2 replication *in vitro* and showed that it can limit, although not completely inhibit, virus replication and disease severity *in vivo* (Fig. 4d, e). This study contributes to the development of new antiviral methods that target proviral factors.

## Methods

### Cells

A549, VeroE6/TMPRSS2 (ref), HuH-7, and 293T cells were cultured in Dulbecco’s modified Eagle’s medium (DMEM) (Thermo Fisher Scientific, Waltham, MA, USA) containing 10% fetal calf serum (FCS) (Thermo Fisher Scientific) and 1% penicillin‒ streptomycin (Nacalai Tesque, Kyoto, Japan). The cells were incubated at 37°C with 5% CO_2_.

### Plasmids

Cas9 and gRNA plasmids were purchased from Addgene (Watertown, MA, USA) (Cas9 vector, 68343; gRNA vector, 67974). These CRISPR-related lentiviruses, including the human v3 library, were produced by cotransfection of 293T cells with the transfer vectors psPAX2 and pMD2.G using Lipofectamine LTX (Thermo Fisher Scientific, Waltham, MA, USA). hACE2 cDNA was inserted between the XhoI and XbaI sites of CSII-CMV-RfA. Lentiviral vectors were rescued through the transfection of these plasmids with pCAG-HIVgp, pCMV-VSVG, and pCMV-Rev into 293T cells using TranIT-293 (TaKaRa, Shiga, Japan) or the ViraPower Lentiviral Expression System (Thermo Fisher Scientific, Waltham, MA, USA).

### Virus preparation

SARS-CoV-2/UT-NCGM02/Human/2020/Tokyo (ref) was propagated in VeroE6/TMPRSS2 cells. For CRISPR‒Cas9 screening, SARS-CoV-2 was purified through ultracentrifugation with 20% sucrose at 27,000 rpm for 2 h at 4°C. The pellets were suspended in Tris-EDTA buffer (pH 8.0) and centrifuged at 3,000 rpm for 10 min at 4°C. The supernatants were stored at −80°C until used for screening. All experiments using SARS-CoV-2 were performed in a biosafety level 3 containment laboratory at Kyoto University, which is approved for use by the Ministry of Agriculture Forestry and Fisheries, Japan. Human coronavirus 229E (HCoV-229E) (ATCC, Manassas, VA, USA) was propagated in HuH-7 cells.

### Generation of Cas9-expressing A549-hACE2 cells

A549 cells were transduced with a lentivirus containing Cas9 and subjected to blasticidin selection (10 μg/ml) (Thermo Fisher Scientific) at 72 h after transduction. A549-Cas9 cells were transduced with a lentivirus containing hACE2 and subjected to Zeocin (200 μg/ml) (InvivoGen, San Diego, CA, USA) at 72 h after transduction. The Cas9 activity of the cells was assessed as described previously (17). Cell lines with Cas9 activity over 98% were used for gRNA library transduction.

### Genome-scale CRISPR/Cas9 screen

The Human CRISPR Library v.3, which targets 18,365 genes with 113,526 sgRNAs, including 1,004 nontargeting control sgRNAs (Addgene, 67989) (18), was used in this study. A total of 3.3 × 10^7^ A549-hACE2-Cas9 cells were transduced with the lentiviral-packaged whole-genome sgRNA library to achieve 30% transduction efficiency (100× library coverage). The cells were subjected to puromycin selection (1 μg/ml) (Thermo Fisher Scientific) at 72 h after transduction. At 10 days after transduction, 3.5× 10^7^ cells were inoculated with purified SARS-CoV-2/UT-NCGM02/Human/2020/Tokyo at a multiplicity of infection (MOI) of 0.3. At 3 days post-infection, cells were harvested and subjected to DNA extraction.

### DNA extraction, gRNA PCR amplification, Illumina sequencing and gRNA counting

Genomic DNA was extracted from cell pellets using the Blood & Cell Culture DNA Maxi Kit (Qiagen, Hilden, Germany) according to the manufacturer’s instructions. PCR amplification, sgRNA sequencing (19-bp single-end sequencing with custom primers) and gRNA counting were performed as described previously (17). The DNB-SEQ platform was used for sgRNA sequencing.

### CRISPR‒Cas9 screen data analysis

To identify hit genes, statistical analysis was performed using MAGeCK v0.5.9.5 (49) by comparing the infected population vs. day 10 population (preinfection), the mock-infected population vs. day 10 population, and the infected and the mock-infected populations.

### Virus titration

The SARS-CoV-2 titre was determined by the median tissue culture infectious dose (TCID_50_) of VeroE6/TMPRSS2. The infectious unit of HCoV-229E was titrated as follows: HuH-7 cells seeded in 96-well plates were inoculated with a serial 10-fold dilution of the virus and incubated for one day. Antigen-positive cells were detected by indirect immunofluorescence assay (IFA) using an anti-HCoV-229E nucleoprotein antibody (Sino Biological, Beijing, China).

### Generation of knockout cells using the CRISPR‒Cas9 system

A549-hACE2-Cas9 cells were transduced with sgRNA targeting 10 genes using lentiviral vectors. The sequences of the sgRNAs used are shown in Supplementary Table 1. The transduced cells were subjected to puromycin selection (1 μg/ml) (Thermo Fisher Scientific) at 72 h after transduction. The knockout efficacy of target genes was assessed by Western blotting.

### Western blotting

Cell lysates were prepared in SDS sample buffer, separated by SDS‒PAGE using e-PAGEL (ATTO Corporation, Tokyo, Japan), and transferred to polyvinylidene difluoride membranes from a Trans-Blot Turbo PVDF Transfer Pack (Bio-Rad, Richmond, USA). The membranes were blocked with 5% skim milk (Wako Pure Chemical Industries) in TBS-T (Tris-buffered saline, 0.1% Tween 20) and incubated with anti-TRIM28 antibody (1:3,000) (sc-515790, Santa Cruz Biotechnology, Dallas, TX, USA), anti-TRIM33 antibody (1:1,000) (ab47062, Abcam, Cambridge, UK), anti-EHMT1 antibody (1:500) (35005S, Cell Signaling Technology, Danvers, MA, USA), anti-EHMT2 antibody (1:1,000) (ab185050, Abcam), anti-VPS35 antibody (1:1,000) (sc-374372, Santa Cruz Biotechnology), anti-SARS-CoV-2 S antibody, or anti-tubulin antibody (1:2,000) (T5168, Merck, Darmstadt, Germany) diluted with Can Get Signal Immunoreaction Enhancer Solution 1 (Toyobo, Osaka, Japan) at 4°C overnight. After washing with TBS-T, the membranes were incubated with a 50,000-fold-diluted HRP-labelled anti-mouse or rabbit IgG antibody (Merck) at room temperature for 2 hours or more. After washing with TBS-T, Clarity Western ECL Substrate (Bio-Rad) was used for detection by chemiluminescence reaction. Bands were detected and photographed by a Fusion Solo instrument (Vilber-Lourmat, Marne-la-Vallée Cedex, France). Band intensities were analysed using ImageJ.

### RT‒qPCR

The virus RNA in the cell or culture supernatant was extracted with nucleospin RNA plus (TaKaRa) and quantified by RT‒qPCR using Primer/Probe N2 (NIHON GENE RESEARCH LABORATORIES, INC. Miyagi, Japan) and One Step PrimeScript™ III RT‒qPCR Mix (TaKaRa) with CFX96 (Bio-Rad, Hercules, CA, U.S.A.) for the N gene. RNA was extracted from sgTRIM28, sgTRIM33, sgEHMT1, and sgEHMT2 cells using nucleospin RNA plus (TaKaRa), and RT‒qPCR was performed using Luna Universal qPCR Master Mix (New England Biolabs, New England Biolabs, Ipswich, MA, USA) and CFX96 (Bio-Rad) to quantify the mRNA levels of IFNb1 and CXCL10. IFNb1-specific primers (forwards, 5’-CGC CTT GGA AGA GTC ACT CA-3’, reverse, 5’-GAA GCC TCA GGT CCC AAT TC-3’) and CXCL10-specific primers (forwards, 5’-GTG GCA TTC AAG GAG TAC CTC-3’, reverse, 5’-GCC TTC GAT TCT GGA TTC AGACA-3’) (50) were used for each.

### Indirect immunofluorescence assay

Cells were fixed with 4% paraformaldehyde (Wako Pure Chemical Industries, Osaka, Japan) at room temperature and treated with PBS containing 0.5% Triton-X 100 and 2% FCS for 15 minutes, followed by incubation with anti-SARS-CoV-2 S antibody (1:1,000) (ab272504, Abcam) for 1 h. After washing, the cells were reacted with a 1000-fold diluted Alexa Fluor rabbit 555-conjugated secondary antibody (Thermo Fisher Scientific), 100 nM Acti-stain 488 Fluorescent Phalloidin (Cytoskeleton, Denver, CO, USA), and 300 nM DAPI (Merck, Darmstadt, Germany) for 1 h. After washing with PBS, the cells were mounted with ProLong™ Diamond Antifade Mountant (Thermo Fisher Scientific). Immunostained cells were observed using a Ti-E inverted microscope with a C1 confocal laser scanning system (Nikon, Tokyo, Japan).

### Electron microscopic analysis

A549-hACE2-Cas9 cells infected with SARS-CoV-2 were fixed with 2.5% glutaraldehyde (TAAB Laboratories Equipment Ltd) and postfixed with 1% osmium tetroxide (TAAB Laboratories Equipment Ltd) at 4°C. After *en bloc* staining with 1% uranyl acetate, the cells were dehydrated with a series of ethanol gradients, followed by propylene oxide, and embedded in Epon 812 resin (TAAB Laboratories Equipment Ltd). The ultrathin sections were stained with 2% uranyl acetate and lead citrate and observed using a Hitachi HT-7700 instrument at 80 kV.

### Entry assay using VSV vector pseudotyped with VSV-G or SARS-CoV-2-S

293T cells were transfected with pCMV-VSV-G (Addgene) expressing VSV-G or pEF4/HisA (Thermo) expressing codon-optimized SARS-CoV-2 spike with a 19 amino acid deletion at the C-terminus using TransIT-293. Twenty-four hours later, G-complemented VSVΔG-GFP (51, 52) was inoculated at an MOI of 0.5. The supernatant was collected 24 hours after inoculation, and debris was removed by centrifugation and stored at −80 degrees. The efficiency of cell entry mediated by each envelope protein was calculated by counting the number of GFP-expressing cells with an Eclipse TE-2000-U inverted microscope (Nikon) after inoculating the generated pseudotyped virus into target cells.

### Animal experiments

SARS-CoV-2 was intranasally inoculated into 4-week-old male Syrian hamsters (Japan SLC Inc, Shizuoka, Japan) at a dose of 10^2^ TCID_50_ per animal using a mixture of 3 anaesthetics. For mock infection, DMEM with 2% FCS was used. UNC0642 at a dose of 5 mg/kg or 0.1% DMSO was administered intraperitoneally once daily starting from one day after infection. Body weight was measured daily from the day of virus inoculation. Four days after infection, the animals were euthanized, and the lungs were removed for virus titre measurement and pathological analysis. The animal experiments were approved by the Animal Experiment Committee at Kyoto University (#A22-8).

### Pathologic assessment

Lungs collected from animals 4 days postinfection were fixed in 10% neutral buffered formalin (Wako), embedded in paraffin (Thermo), sectioned, and stained with haematoxylin (Merck) and eosin (Wako). Pathological features in airways, interstitium, vessels, alveolar spaces, oedema, and alveolar epithelial regeneration were scored as previously reported (38, 39), with a score of 0 (no lesions), 1 (mild), 2 (moderate), and 3 (severe).

## Supplementary Table

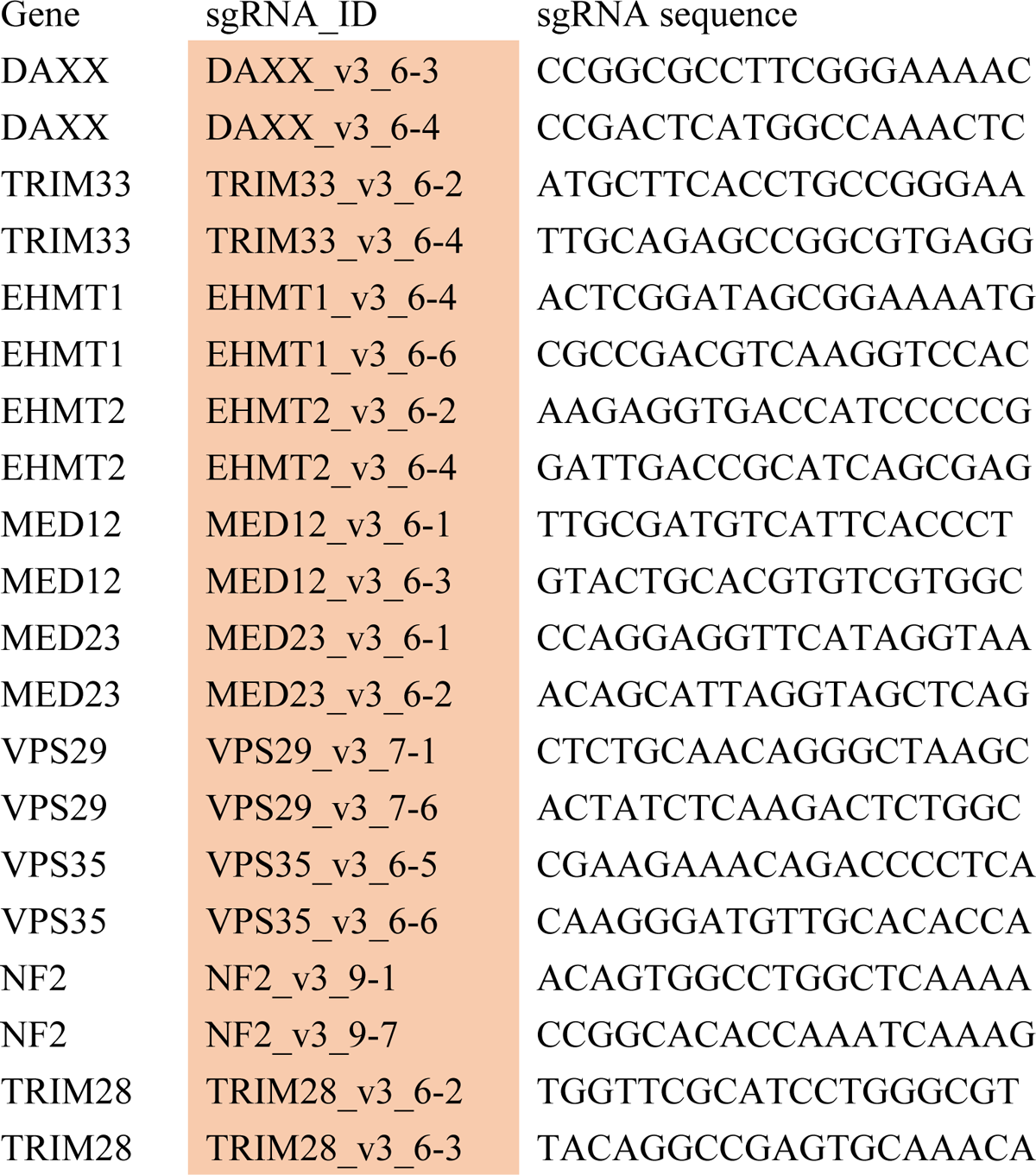

---

**Supplementary Data 1.** CRISPR‒Cas9 screening data analysis. (a) Scatterplot of gRNA counts in gCTRL (n=1004, highlighted in orange) cells to confirm the quality of screening. Comparison between preinfection versus mock and preinfection versus infected. (b) Volcano plot of gRNA change compared between mock and infected cells.

**Supplementary Data 2.** Knockout efficiency of selected genes in A549-hACE2-Cas9 cells. **(a)(b)** Western blotting analysis of selected genes, TRIM28 and TRIM33 (a) and EHMT1 and EHMT2 (b). The expression levels were evaluated by the band intensity, which was normalized to that of α-tubulin. Data are presented as the mean ± the SD of three independent experiments. Dunnett’s multiple-comparison test. *, *p* < 0.05; **, *p* < 0.01.

**Supplementary Data 3.** (a) Cytotoxicity of UNC0642. Cell proliferation under UNC0642 treatment was assessed using a WST-1 assay (TaKaRa). **(b)** Inhibitory effect of remdesivir on SARS-CoV-2 infection. Statistical analysis was performed using Dunnett’s multiple-comparison test, and every point did not show significance.

**Supplementary Data 4.** Pathological scoring in airways, interstitium, vessels, alveolar spaces, oedema, and alveolar epithelial regeneration. Violin plot of inflammation scores in the lungs of hamsters 4 days post-infection.

## Acknowledgements

This work was supported in part by the Japan Society for the Promotion of Science (JSPS) KAKENHI grant numbers JP19K22530 (KT), JP20H00662 (KT), JP21K19909 (KT) and 18K05991 (AM); National BioResource Project of the Ministry of Education, Culture, Sports, Science and Technology (MEXT) KAKENHI grant numbers 19H04834 (AM); Japan Agency for Medical Research and Development (AMED) grant number JP20wm0325011h0001 (AM&KY); Japan Science and Technology Agency (JST) Start-ups from Advanced Research and Technology (START) Project Promotion Type (Commercialization Support), Grant Number JPMJST2113 (KT); JSPS Core-to-Core Program JPJSCCA20190008 (TN&KT); 2021 Kaketsuken Research grant (KT); JST Core Research for Evolutional Science and Technology grant number JPMJCR20HA (TN); and the Joint Usage/Research Center Program on Institute for Life and Medical Sciences, Kyoto University. CSII-CMV-RfA, pCAG-HIVgp, pCMV-VSVG, and pCMV-Rev were provided by the RIKEN BRC through the National BioResource Project of the MEXT/AMED, Japan.

## Author contributions

A.M. designed and managed the overall project. M.S., Y.M., Y.T., M.I., R.K., K.Y., K.T., and A.M. performed and analysed the *in vitro* experiments. Y.T. and T.N. performed electron microscope analysis. M.S. and A.M. conducted animal experiments. N.A. and Y.K. performed the pathological assessment.

