## Supplementary figures and images for "Genome-scale CRISPR‒Cas9 screen identifies novel host factors as potential therapeutic targets for SARS-CoV-2 infection"

### Supplemental Data

a

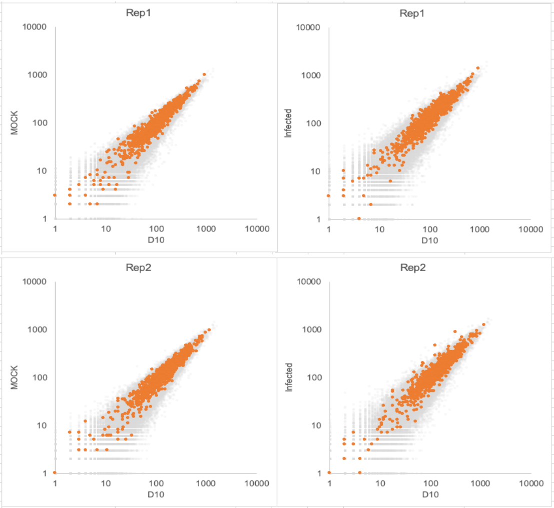


b

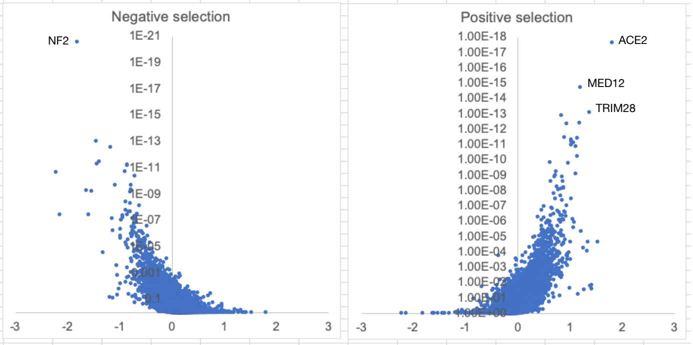


Supplementary Data 1

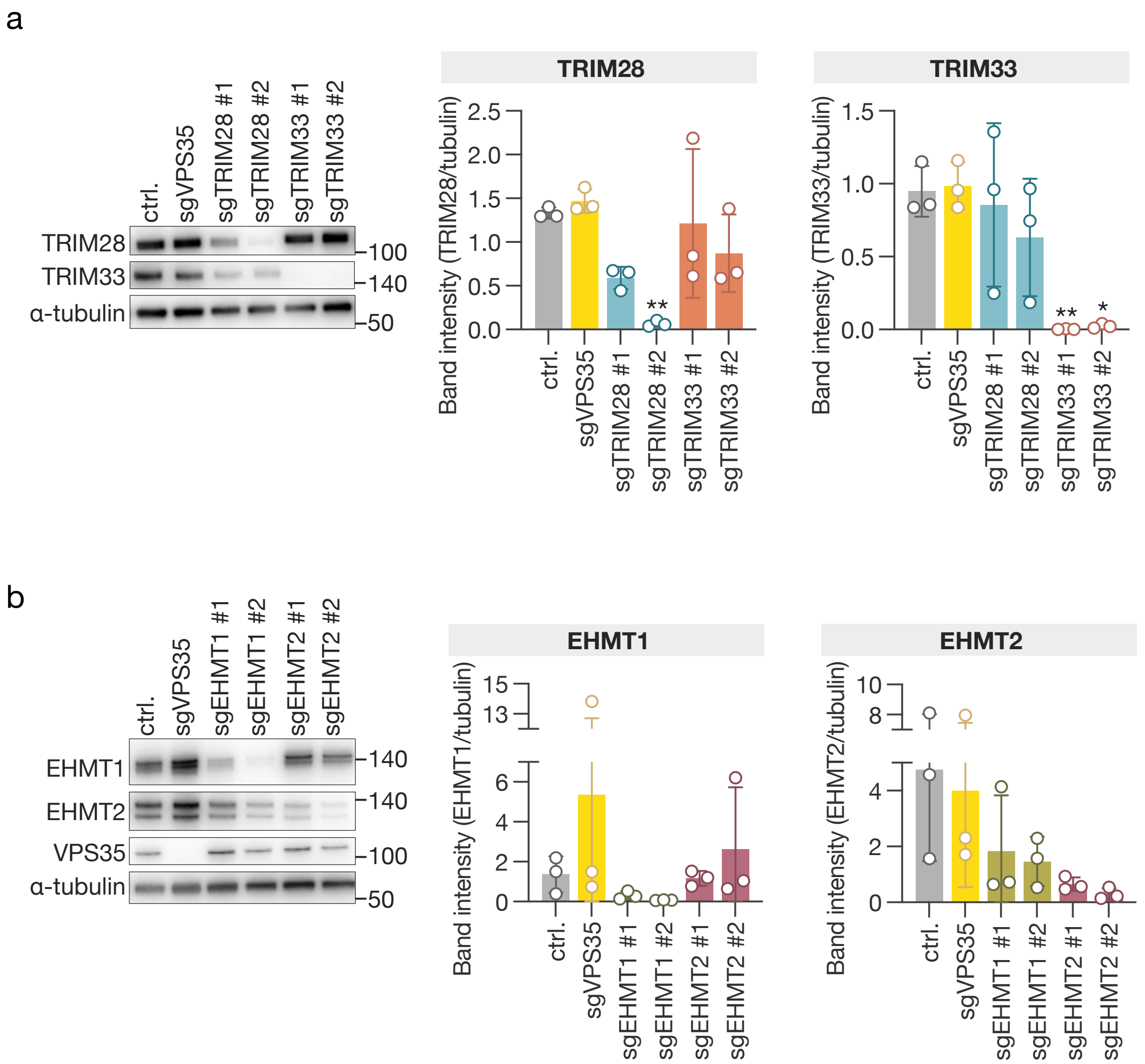


Supplementary Data 2

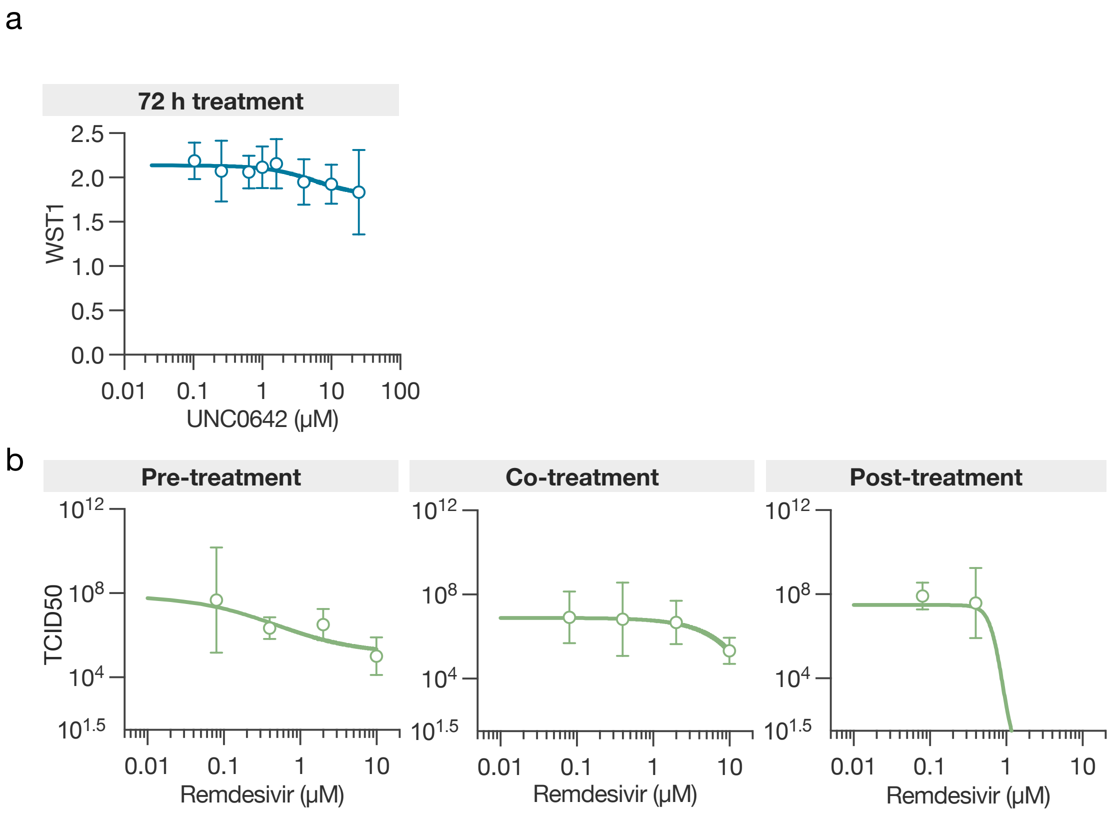


Supplementary Data 3

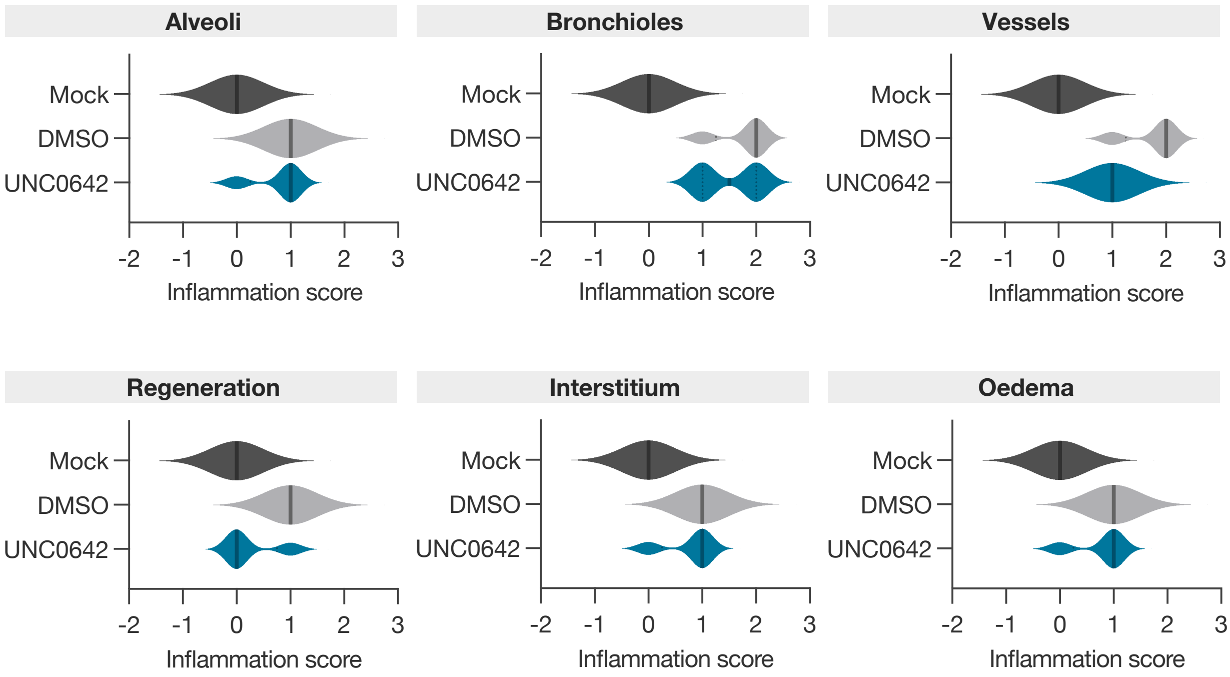


Supplementary Data 4
